# The functional role of locus coeruleus microglia in the female stress response

**DOI:** 10.1101/2024.01.10.575076

**Authors:** Cora E. Smiley, Brittany S. Pate, Samantha J. Bouknight, Evelynn N. Harrington, Aaron M. Jasnow, Susan K. Wood

## Abstract

Neuropsychiatric disorders that result from stress exposure are highly associated with central inflammation. Our previous work established that females selectively exhibit heightened proinflammatory cytokine production within the noradrenergic locus coeruleus (LC) along with a hypervigilant behavioral phenotype in response to witnessing social stress, and ablation of microglia using pharmacological techniques prevents this behavioral response. These studies were designed to further investigate the impact of stress-induced neuroimmune signaling on the long-term behavioral and neuronal consequences of social stress exposure in females using chemogenetics. We first characterized the use of an AAV-CD68-G_i_-DREADD virus targeted to microglia within the LC and confirmed viral transduction, selectivity, and efficacy. Clozapine-n-oxide (CNO) was used for the suppression of microglial reactivity during acute and chronic exposure to vicarious/witness social defeat in female rats. Chemogenetic-mediated inhibition of microglial reactivity during stress blunted the neuroimmune response to stress and prevented both acute and long-term hypervigilant behavioral responses. Further, a history of microglial suppression during stress prevented the heightened LC activity typically observed in response to stress cues. These studies are among the first to use a chemogenetic approach to inhibit microglia within the female brain *in vivo* and establish LC inflammation as a key mechanism underlying the behavioral and neuronal responses to social stress in females.

## Introduction

Stress exposure is a risk factor for many physiological and neuropsychiatric disorders including post-traumatic stress disorder (PTSD) and substance use disorder (Cohen et al., 2007; Liu et al., 2017). While up to 90% of the general population reports experiencing a highly stressful or traumatic event (Benjet et al., 2016), around 15% of these people will go on to develop conditions such as PTSD (Davidson, 2000). Interestingly, while males report greater levels of overall trauma exposure, females develop PTSD at twice the rate of males (Frans et al., 2005). Further, there are different symptom patterns between the sexes with the hyperarousal and hypervigilance aspects of PTSD symptomology (Miao et al., 2018) observed to a higher degree in females (Bangasser & Valentino, 2014; Irish et al., 2011). These symptoms are marked by an overall higher arousal state, where patients report feeling “on edge”, are constantly attending to stimuli even in safe and neutral environments, and display an exaggerated startle response (Kimble et al., 2014; Sadeghi et al., 2022). Additionally, contextual and environmental cues that are reminiscent of the initial traumatic event are capable of inducing a hypervigilant state (Liberzon & Abelson, 2016). Early emergence of hyperarousal and hypervigilance are also associated with less symptom improvement over time and worse treatment outcomes when compared to traumatized individuals that did not exhibit this symptom profile (Marshall et al., 2006; Schell et al., 2004). Thus, there may be inherent neurobiological factors that promote hypervigilance and are responsible for the 2-3x greater risk of developing stress-related disorders in females (Bangasser & Valentino, 2014).

One brain region highly involved in stress responses and hypervigilance is the noradrenergic locus coeruleus (LC) (Bangasser & Valentino, 2014; Borodovitsyna et al., 2020; Giustino et al., 2020; C. Naegeli et al., 2018; O’Donnell et al., 2004). Exposure to stress increases activity within this region, and LC hyperactivity is evident in patients with PTSD (Krystal et al., 2018) and directly promotes hypervigilance (McCall et al., 2015; C. Naegeli et al., 2018). As the main noradrenergic center of the brain, increased activity in the LC promotes norepinephrine (NE) release in multiple downstream brain regions that mediate stress-related behaviors (Borodovitsyna et al., 2020; McCall et al., 2017). Therefore, further research into the underlying mechanisms responsible for stress-induced increases in LC-NE activity is essential for understanding the role of this system in the hypervigilance-related behavioral dysfunction observed following traumatic stress exposure. Importantly, the LC has been shown to be highly regulated by neuroimmune signaling, with proinflammatory cytokines selectively administered within this region leading to increased neuronal firing (Borsody & Weiss, 2002; Borsody & Weiss, 2004). This function of neuroimmune signaling may serve as an underlying factor responsible for the increases in LC activity observed in response to stress. In accordance with this hypothesis, PTSD, among other highly prevalent stress-induced neuropsychiatric disorders, is associated with increases in inflammatory factors in the brain and periphery (Hänsel et al., 2010; Ravi et al., 2021; Smiley & Wood, 2022). Further, neuroimmune signaling has been associated with psychiatric dysfunction following stress in rodent models (Finnell et al., 2018; Pate et al., 2023) and in clinical populations (Engler et al., 2016), especially in females (Bekhbat & Neigh, 2018; Levesque et al., 2023). Specifically, lipopolysaccharide (LPS) injections evoke greater inflammatory, noradrenergic, and cortisol responses in women which were associated with worsening mood symptoms that were not observed in men (Engler et al., 2016; Moieni et al., 2015). Overall, these data support the hypothesis that the predisposition to stress-related neuropsychiatric disorders in females may be due to an underlying sensitivity to the neuroimmune impact of stress.

Despite accumulating evidence that suggests a role for neuroimmune signaling in heightened stress responding in females, studies examining a specific role for microglia are sparse, largely due to a lack of effective tools in the field. Our previous studies using clodronate, a compound designed to pharmacologically ablate microglial cells, support the hypothesis that microglia within the LC play a major role in the female stress response (Pate et al., 2023). Therefore, by utilizing chemogenetics to transiently inhibit microglial activation during social stress exposure, these studies were designed to further explore the role of neuroimmune signaling in the acute and long-term behavioral consequences associated with hypervigilance in females. Further, the impact of prior intra-LC microglial inhibition on neuronal activity was assessed in response to stress cues to determine how microglia shape future stress context-dependent neuronal activation. DREADD viruses designed to chemogenetically inhibit microglia have been previously characterized for use in the spinal cord (Grace et al., 2016; Grace et al., 2018) and, through G-protein coupled receptors (GPCRs), can prevent microglial activation and subsequent release of proinflammatory cytokines (Gu et al., 2021; Haque et al., 2018; Kettenmann et al., 2011; Mizoguchi & Monji, 2017). Initial reports using these viral constructs conducted *in vitro* experiments in which microglial cells were cultured and transfected with either hM3D_q_ or hM3D_i_ viral constructs before treatment with LPS and CNO. These studies determined that G_i_-mediated suppression of LPS-induced microglial activity using CNO reduced the RNA levels of eight major inflammatory signaling molecules, including the proinflammatory cytokine IL-1β, while these factors were increased when G_q_ receptors were stimulated with CNO (Grace et al., 2018). These effects were then replicated *in vivo* using rat models with DREADD administration in the spinal cord (Grace et al., 2016; Grace et al., 2018). However, species specific differences in the immune system may have previously limited the use of this tool in preclinical experiments. The genetic profile of microglia cells and level of cytokine release in response to inflammatory insults varies between murine species (Lam et al., 2017; Lively et al., 2018). Specifically, rats exhibit six times greater levels of CD68 in microglial cells when compared to mice, which may lead to a better transduction efficiency of viruses with a CD68 promoter when applied to rats compared to mice (Lam et al., 2017). Therefore, when testing novel immune strategies for the translational treatment of neuropsychiatric disorders and new applications of preclinical techniques targeting immune cells, it is important to distinguish between results that have been achieved in different species. Thus, these experiments applied chemogenetic techniques to rat models to modulate microglial activity during acute and repeated psychosocial stress to assess the short- and long-term behavioral impact of stress-induced neuroimmune signaling in the LC of female rats.

## Materials and Methods

### Animals

Female Sprague-Dawley rats (∼200 g and 9 weeks on arrival, witnesses/controls) and male Sprague-Dawley rats (∼250 g on arrival, intruders) were obtained from Charles River (Durham, NC) while male Long-Evans retired breeders (600-800 g, residents) were obtained from Envigo (Dublin, VA). All rats were housed individually in standard polycarbonate cages. Female rats that underwent stress, and all males, were housed in a separate but adjacent room from control females. Rats were all maintained on a 12-hr light/dark cycle with the light cycle initiated at 0700 and received *ad libitum* access to food and water. Further, all cages, including the home cage and those used during behavioral testing, contained Teklad sani-chip bedding (Envigo). These studies were approved by the University of South Carolina’s Institutional Animal Care and Use Committee and maintained adherence to the National Research Council’s Guide for the Care and Use of Laboratory Animals.

There were four main experimental subsets completed to characterize the use of chemogenetic techniques for the site-specific manipulation of microglial activity in the rat brain and determine the role of LC microglia in the behavioral and neuronal responses to social stressors in females. Experiment #1 used 8 rats who had the pAAV1 CD68-hM4D(G_i_)-mCherry virus infused into the LC to determine the cell type selectivity of viral transduction into microglia, astrocytes, or neurons. Experiment #2A used 22 rats (n = 7-8/group, CON+VEH vs. WS+VEH vs. WS+CNO) to determine the impact of viral activation with CNO on microglial-mediated cytokine production in response to stress. Experiment #2B used 36 rats (n = 6/group for behavioral analysis, 4/group for microglial analysis) to establish the behavioral impact of chemogenetic suppression of microglial activity on stress-induced behavioral responding and LC microglial morphology. A separate group of rats (Experiment #2B subset with no surgical manipulation) were exposed to acute stress and assessed for behavioral responses and microglial ramification characteristics in the absence of viral infusion and in the presence of CNO treatment to determine any off-target effects of CNO (n = 28, 7/group for behavioral analysis, a subset of 4/group for microglial analysis). Experiment #3 used 16 rats (n = 4/group, VEH or LPS + VEH or CNO) to assess the impact of CNO-mediated microglial suppression in response to LPS on microglial morphology in the LC. Experiment #4 used 34 rats (n = 8-9/group) to assess the behavioral impact of acute and repeated vicarious/witness defeat stress in the presence and absence of DREADD-mediated suppression of microglial responding and resulting behavioral and neuronal responses to stress context presentation. Experimental design details are presented in **Figure 1**.

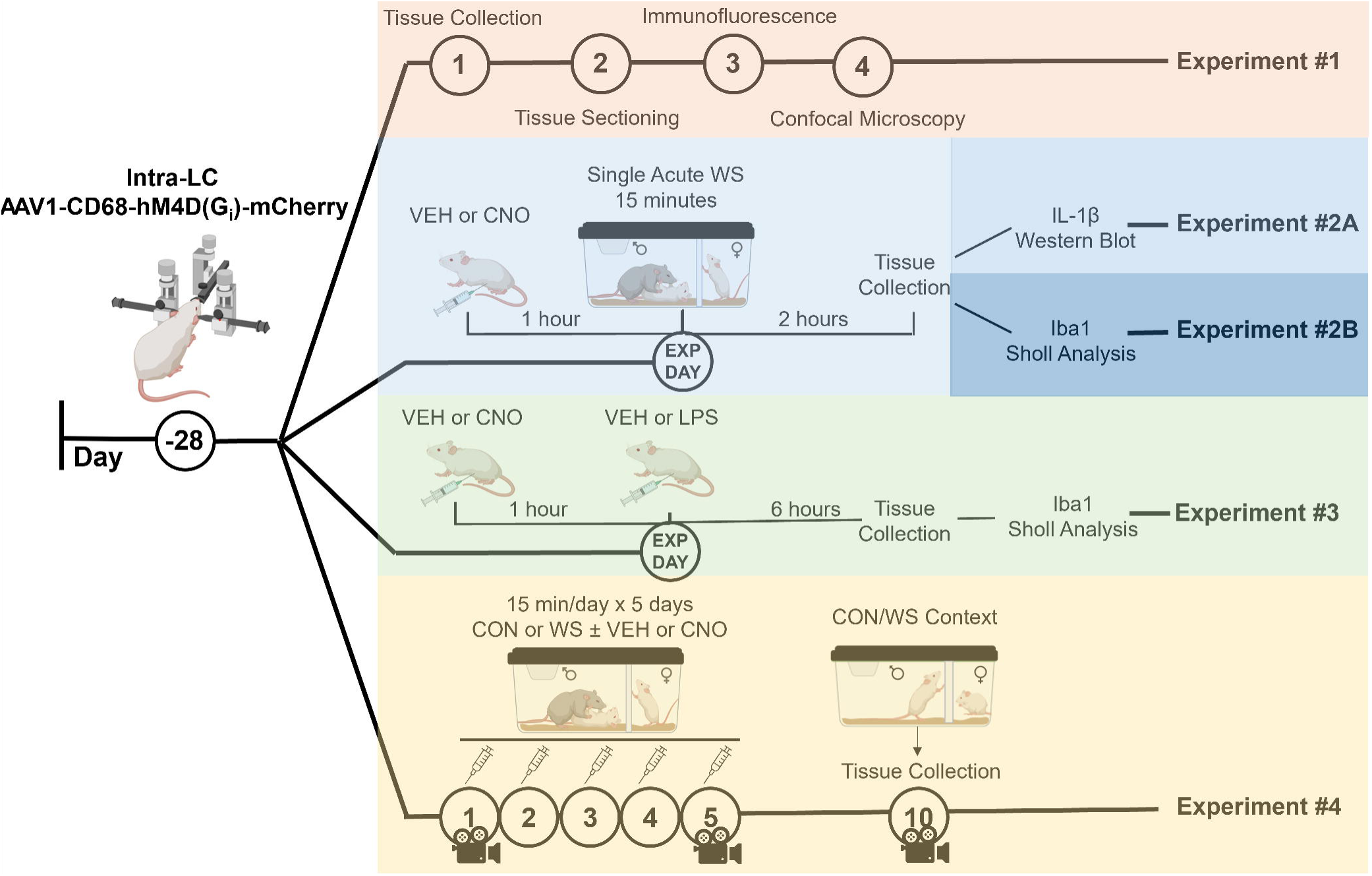

### Surgical Procedures and Viral Infusion

All experiments, excluding the Experiment #2B subset designed to assess off-target effects of CNO, commenced with stereotaxic surgery to infuse a pAAV1 CD68-hM4D(G_i_)-mCherry virus (Addgene viral prep #75033-AAV1 from Bryan Roth, Addgene, Watertown, MA) into the LC. The CD68 promoter allows selective targeting of microglial cells, and this virus has been previously characterized for microglial localization and inhibition in the spinal cord (Grace et al., 2016; Grace et al., 2018; Yi et al., 2021). While typically an AAV lacking the G_i_ component is utilized as the control condition for chemogenetic experiments, these studies were limited by the lack of a commercially available pAAV1 CD68-mCherry virus and, therefore, all rats were injected with the active virus and systemic vehicle injections were used in parallel with CNO administration to control for viral activation.

Rats were anesthetized using isoflurane (5% for induction, 2-3% maintenance) and placed into a stereotaxic apparatus. During surgery, electrophysiological recordings were used for precise identification of the LC to individually localize viral injection. The nose was angled down 10° and, starting at the following coordinates, in mm from bregma and the surface of the dura, A/P: −4.0; M/L: ± 1.2, D/V: −4.5, a tungsten coated electrode was lowered through the tissue while an oscilloscope relayed neuronal firing patterns. Once the LC was located, these coordinates were used to lower a microsyringe (Hamilton, Franklin, MA) to inject 500 nL of the pAAV CD68-hM4D(G_i_)-mCherry virus (100 nL/minute) and the needle was left in place for at least 5 minutes following termination of the injection. These same coordinates were used for the contralateral LC to achieve bilateral infusion of the virus. Saline (10 mL/kg, s.c.) and flunazine (0.25 mg/kg, s.c.) were administered on the day of and day following surgery along with supplemental nutrition (Bacon Softies, Bio-Serv) as needed. Rats were given at least 28 days prior to stress to allow for viral expression and, during this recovery period, rats were given i.p. saline injections three times per week for habituation prior to drug delivery. Viral localization to the LC was confirmed with microscopic visualization of 30-micron tissue slices and injections sites for each treatment group are displayed in the supplement (**Figure S1**). Although electrophysiological confirmation of localization during surgery greatly reduces the number of off-target injections, one animal was discovered to have unilateral viral placement and was eliminated from analysis. Further, a subset of surgically naïve animals that received CNO in the absence of viral infusion was included to confirm that CNO alone did not impact microglial reactivity or stress-related behaviors.

### Drug Information

Water soluble clozapine-n-oxide (CNO) (HelloBio, Princeton, MA) was dissolved in sterile saline (0.9%) and rats were administered CNO (1 mg/kg) or vehicle (0.9% saline) intraperitoneally (i.p.) one hour before the start of stress either once (acute stress) or for five days in a row (repeated stress). Lipopolysaccharide was diluted in saline to 3 µg/µL and was administered at a dose of 30 µg/kg one hour following CNO in Experiment #3.

### Behavioral Measurements

#### Witness Stress or Control Handling

Female rats in all experiments were exposed to witness stress paradigms (WS) as previously described (Finnell et al., 2018; Pate et al., 2023; Smiley et al., 2023). Briefly, a clear perforated Plexiglas partition was placed into the cage of a male Long-Evans retired breeder (resident) to allow for the transfer of visual, olfactory, and auditory cues without physical access (described in detail in (Finnell et al., 2018)). The female witness was placed behind the partition followed immediately by the male intruder into the side containing the resident, and witnesses observed the aggressive social defeat encounter for 15 minutes. Female rats were either exposed to a single acute stressor (Experiment #2) or this paradigm was completed once a day for a total of five days for those that received repeated stress (Experiment #4), and CNO or vehicle was administered one hour prior to the start of each stress exposure. Witnesses were paired with the same male intruder rat for daily WS, but the resident was novel each day to maintain attack behaviors. Control rats (CON) were briefly handled and returned to their home cage where behaviors were recorded. Previous experiments from our lab established that females exposed to the resident’s cage behind the partition with the intruder on the opposite side (Finnell et al., 2018) or in absence of the intruder (Smiley et al., 2023), do not exhibit behavior that differs from rats handled and returned to their home cage, thus, this procedure was utilized as the control. Sessions were video recorded on the first and fifth days of exposure to allow for retrospective manual scoring of stress-evoked behaviors by an experimenter blinded to the treatment groups. Briefly, behaviors including burying, freezing, and rearing were quantified according to the following attributes and our previous publications (Finnell et al., 2018; Pate et al., 2023; Smiley et al., 2023). Burying was counted when the female rat exhibited spontaneous and rapid manipulation of the bedding. Freezing consisted of complete immobility with the exception of respiratory movement or vocalization. Rearing was displayed when the animal raised their forepaws off of the bedding, with or without extension of the hindlimbs. The latency to begin each behavior along with the total duration was quantified and analyzed.

#### Control or Witness Stress Context Exposure (CON/WS CXT)

WS CXT consisted of placing the female rat behind the partition in the residents’ cage with their paired intruder on the opposite side but in the absence of the resident to expose the females to the sensory cues and environment in which they originally experienced social stress. Previous studies from our lab have determined that a history of stress exposure leads to increased burying in the WS CXT and this paradigm serves as a method of translationally measuring hypervigilant behaviors (Smiley et al., 2023). Behaviors were scored identically to those assessed during WS (described above) and were compared to CON rats that were handled and recorded in their home cages.

#### Euthanasia and Tissue Collection

All euthanasia procedures followed the guidelines set forth by the American Veterinary Medical Association (AVMA) and sufficient loss of consciousness was induced by isoflurane prior to any manipulation to ensure minimal pain and distress. Following deep anesthetization with isoflurane, confirmed by a lack of reflexive responding to a toe pinch, rats were either transcardially perfused with 0.1M phosphate buffered saline (PBS) followed by 4% paraformaldehyde (4% PFA prepared in 0.1M PBS) or rapidly decapitated. Rats in Experiment #1 were perfused at rest approximately four weeks after viral infusion for the immunofluorescent assessment of virus cell type specificity. For Experiment #2A, rats were euthanized by rapid decapitation and fresh tissue was collected following acute stress + VEH or CNO for Western Blot analysis of proinflammatory cytokine expression. Experiments #2B and #3 concluded with the collection of perfused brains for immunohistochemical staining of microglial morphology. Rats in Experiment #4 were perfused after stress context presentation and tissue was collected for immunohistochemical assessment of LC neuronal activity. Perfused brains were extracted, blocked into anterior and posterior sections, post-fixed in 4% PFA for at least 2 hours, and saturated with 20% sucrose + 1% azide upon which they were flash frozen using ice cold isopentane and stored at −80°C until further analysis while non-perfused brains were flash frozen prior to dissection.

### Brain Analyses

Perfused brains were sliced at 30 microns on a cryostat (Leica) and slices from the LC were collected and stored in cryoprotectant at −20°C prior to immunohistochemical analysis while non-perfused brains were micropunched for discrete brain regions. Sections were analyzed for mCherry localization to the LC and selectivity for microglial cells, microglial morphology with Iba1 staining (Kedia & Chattarji, 2014), or LC neuronal activation with colabeled cfos and tyrosine hydroxylase (TH) used to label neurons within the LC (Fan et al., 2014). Tissue punches were assessed via Western Blot for the effects of microglial inhibition on cytokine production as a result of stress. Visual confirmation of mCherry and Iba1 colabeling within the LC was completed with confocal microscopy (Leica Stellaris) to ensure that mCherry was selectively localized to microglial cells. Microglial cell body size and ramification state were assessed via tracing using Neurolucida software and quantified with Neurolucida Explorer. Tissue assessments were completed following the protocols outlined below.

#### Immunohistochemistry

Tissue slices were washed 4×10 minutes in 0.1M PBS, immersed for 30 minutes in 0.9% hydrogen peroxide + 4% Triton-X 100 (TX) in 0.1M PBS, and incubated in 1° antibodies overnight at room temperature in antibody buffer consisting of 1% normal donkey serum (NDS) + 0.25% TX in 0.1M PBS. Antibodies were used at the following concentrations: anti-Iba1 1:1000 (rabbit, FujiFilm Wako Chemicals #019-19741), anti-cfos 1:500 (rabbit, Millipore #ABE-457), and anti-TH 1:2500 (mouse, Millipore #MAB-5280). The following day, tissue slices were washed 3×10 minutes in 0.1M PBS prior to a 2-hour incubation at room temperature in secondary antibody (diluted in antibody buffer) that corresponds with the species of the primary: biotinylated anti-rabbit 1:200 (Vector Labs #BA-1000) or biotinylated anti-mouse 1:1000 (abcam #ab208001). Tissue was then washed in PBS (3×10 minutes) and incubated in avidin biotin complex (Vectastain Elite ABC Kit, Vector Laboratories) diluted in 1% NDS in 0.1M PBS for 1 hour. Three subsequent washes (10 minutes each in 0.1M PBS) occurred prior to staining in diaminobenzidine (DAB) or metal enhanced DAB (Vector Laboratories) for double labeling experiments. The DAB development times were optimized for each antibody, with 1 minute and 30 seconds for Iba1, 2 minutes for cfos, and 15 seconds for TH. Tissue was placed in a final set of 3×10 minute washes before mounting on slides (SuperFrost). Once dried, slides were dehydrated in increasing concentrations of ethanol (70%, 95%, 100% x2, each for 1 minute) and placed into HistoClear (2 and 5 minutes) prior to coverslipping with Permount (Fisher Scientific). Slides were imaged using a Microbrightfield microscope and traced by Neurolucida software (MBF Biosciences) for quantification of microglial morphological characteristics or a Nikon E600 microscope and counted for cfos/TH+ cells in Image J (NIH) by an experimenter blinded to the treatment conditions.

#### Immunofluorescence (IF)

IF techniques were improved by utilizing antibody signaling enhancing buffers to increase the permeabilization of cells (Piña et al., 2022). Brain slices were washed 4×10 minutes in 0.5% Tween-20 in 0.1M PBS and incubated for 30 minutes in blocking buffer consisting of 2% NDS, 50 mM glycine, 0.05% Tween-20, 0.1% TX, and 0.01% bovine serum albumin (BSA) in 0.1M PBS. Sections were then placed into primary antibody overnight at 4°C in antibody buffer containing 10 mM glycine, 0.05% Tween-20, 0.1% TX, and 0.1% hydrogen peroxide in 0.1M PBS along with either anti-Iba1 1:1000 (rabbit, FujiFilm Wako Chemicals #019-19741), anti-NeuN 1:500 (mouse, Millipore #MAB-377), or anti-GFAP 1:500 (rabbit, Cell Signaling #12389T) primary antibody. Subsequently, tissue was rinsed in 0.5% Tween-20 in 0.1M PBS (3×10 minutes) and incubated in the secondary antibody (anti-rabbit Alexa-Fluor 488, 1:500, abcam #150073 or anti-mouse Alexa-Fluor 488, 1:500, abcam #ab1501113) diluted in 0.1% Tween-20 in 0.1M PBS. Slices were then washed 3×10 minutes in 0.1M PBS, mounted onto slides (SuperFrost), and coverslipped with Prolong Diamond Media (Thermo Fisher). Slides were imaged on a confocal microscope (Leica Stellaris) for localization of viral-mediated mCherry with Iba1+ microglial cell staining.

#### Western Blot Analysis

LC tissue punches were collected for Western Blot analysis of IL-1β protein concentration. Flash frozen brains were sliced on a cryostat (Leica) starting at the brain stem, until reaching the most posterior point of the LC, at which time a tissue biopsy punch (1 mm in diameter, 1 mm depth) was used to selectively dissect out the LC. Post punch slices were collected and stained with Neutral Red to confirm that all LC cells were removed (supplemental **Figure S2**). Tissue was homogenized with 200 µL of lab-made lysis buffer (37 mM NaCl, 20 mM Tris, 1% Igepal, 10% glycerol, and 1X Halt Protease Inhibitor Cocktail [Thermo Scientific]) and 100 mg of zirconium oxide beads (Next Advance, Inc., Troy, NY) in a Bullet Blender (3 minutes at speed 8) followed by centrifugation at 14000 rcf for 15 minutes at 4°C. Supernatant was collected and a 12 µL aliquot was diluted 1:4 with 0.1 PBS + 1% azide for protein concentration analysis using a Pierce Bicinchoninic Acid (BCA) assay (Thermo Scientific, Rockford, IL) per the manufacturers protocol. Samples containing 20 µg of protein were aliquoted for Western Blotting. 6X sample buffer + β-mercaptoethanol was added at a 1:6 ratio with the LC homogenate and heated at 75°C for 5 minutes. LC homogenate was loaded onto mini-protean gels (Bio-Rad, Hercules, CA), placed into electrophoresis chambers filled with 1X running buffer (Bio-Rad), and run at 135 V for 1 hour. Proteins were transferred from gels onto PVDF membranes in 1X transfer buffer (Bio-Rad) and run at 100 V for 90 minutes. Membranes were blocked in a solution containing a 1:1 ratio of 0.1M PBS:Fluorescent Blocking Buffer (Millipore Sigma, Burlington, MA) for 1 hour prior to incubation in primary antibodies overnight a 4°C. The antibodies for IL-1β (1:200, goat, R & D Systems #AF-501-NA) and the housekeeping protein GapDH (1:1000, mouse, Santa Cruz #sc-365062) were diluted in blocking buffer + 0.2% Tween-20. Membranes were washed in 1X TBS + 1% Tween-20 before incubation in secondary antibodies IR-Dye anti-rabbit 800 nm (LI-Cor #926–32213) and IR-Dye anti-mouse 680 nm (LI-Cor #962–68072) diluted 1:20,000 in blocking buffer + 0.2% Tween-20 + 0.1% sodium dodecyl sulfate (SDS) for 1 hour. Membranes were washed (3×10 minutes in 1X TBS + 1% Tween-20), imaged on a LI-Cor Odyssey scanner (LI-Cor Biotechnology, Lincoln, NE), and protein expression was quantified using LI-Cor Image Studios Software with normalization to GapDH. Full Western Blot images are provided within the supplement (supplementary **Figure S3**).

#### Analysis of Microglial Ramification Characteristics

Microglial morphology was quantified using Neurolucida Explorer software to trace individual microglia cell bodies and their projections. LC tissue sections labeled with Iba1 and stained with DAB were imaged at 100x magnification on a Microbrightfield microscope equipped with Neurolucida software (MBF Biosciences) for cellular tracing and reconstruction. Four rats from each treatment group were randomly chosen for morphological analysis with four LC tissue sections assessed per animal and five cells traced per section. For each rat, the data for each tissue section was averaged to allow for a n = 20 cells/animal and n = 4 animals per group. Microglial reconstructions from Neurolucida were imported into Neurolucida Explorer software for automated quantification of key morphological features including the cell body area, the number of projection endpoints, the longest projection, the total length of all projections, and the sholl curve which was established based on the number of projections intersecting increasing diameters from the cell body. Intersections were measured at 5-micron increments starting 2.5 microns away from the cell body center. Detailed images depicting the imaging, tracing, and reconstruction of microglial cells are displayed in **Figure 4A & B** and **Figure 5A & B**.

#### Statistical Analysis

All analyses were completed using GraphPad Prism 9 (GraphPad Software, La Jolla, CA). Data were analyzed using either an unpaired t-test, to determine differences between stressed animals treated with either VEH or CNO, or two-way ANOVA with Tukey post-hoc analysis, to compare data in experiments with a 2×2 design assessing CON vs. WS and VEH vs. CNO. Data are presented as mean ± standard error (SEM) with an α = 0.05. Significant main effects are reported in the text with significant post hocs reported on the figures.

## Results

### AAV-CD68 is selectively expressed in microglia

Experiment #1 used brain tissue collected from female rats ∼28 days following intra-LC viral infusion. The CD68 promoter allows selective insertion of G_i_ receptors onto microglial cells, and this virus has been previously characterized for microglial localization and inhibition in the spinal cord (Grace et al., 2018; Yi et al., 2021). LC slices were collected and stained for the microglial marker Iba1, astrocyte marker GFAP, or neuronal marker NeuN. Immunofluorescent labeling of Iba1 was visualized along with mCherry to confirm that virus was solely incorporated into microglial cells. Confocal analysis of these tissue sections revealed microglial cell type specificity of viral expression with selective colocalization of mCherry with Iba1 (**Figure 2 Row 1**) and mCherry overlap absent in astrocytes (**Figure 2 Row 2**) and neurons (**Figure 2 Row 3**).

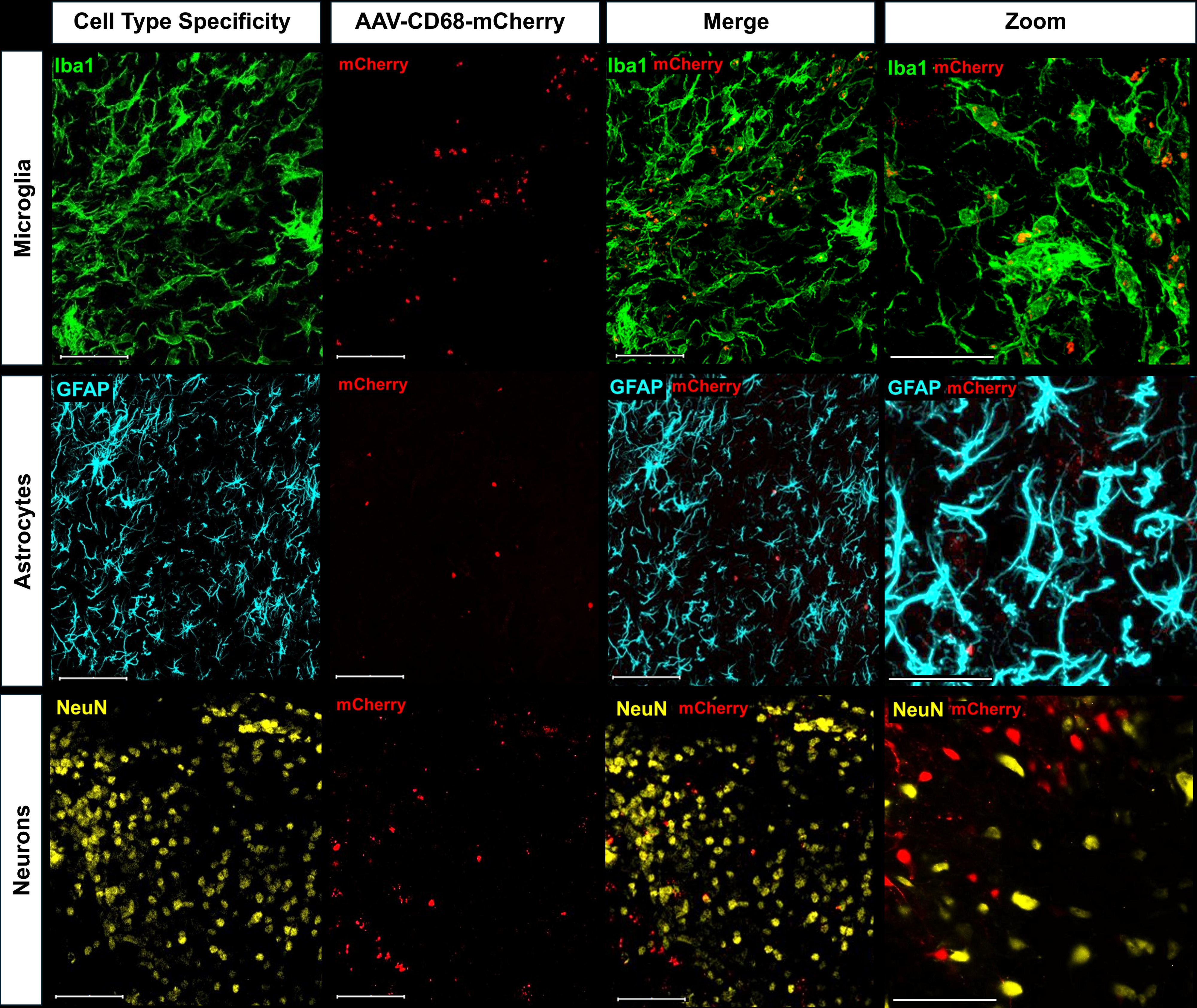

### Chemogenetic inhibition leads to a functional reduction in stress-evoked cytokine production

Four weeks after intra-LC DREADD infusion, rats in Experiment #2A were treated with VEH or CNO one hour prior to a single acute WS exposure followed by brain tissue collection. Behavior was assessed during stress, and LC micropunches were collected, homogenized, and assessed for IL-1β cytokine protein content via Western Blot. In accordance with our previous studies (Finnell et al., 2018; Pate et al., 2023; Smiley et al., 2023), WS exposed females treated with vehicle exhibited significantly higher levels of burying when compared to controls (**Figure 3A**, *p* = 0.0001). However, if females were treated with CNO one hour prior to WS, this effect was diminished and this group displayed significantly less burying in response to the stressor that was not different than controls (**Figure 3A**, one way ANOVA *F_2,19_* = 14.44, *p* = 0.0002, WS+VEH vs. WS+CNO p = 0.008, WS+CNO vs. CON p =0.1184). One animal in the WS+VEH group was eliminated from analysis due to sickness behavior present during the video recording. These behavioral results were paralleled by molecular analysis of the proinflammatory cytokine IL-1β in the LC. Exposure to acute WS evoked an increase in levels of IL-1β within the LC (p = 0.0214) an effect that was diminished by DREADD activation with CNO, with CNO treated stressed female exhibiting IL-1β levels comparable to controls (**Figure 3B**, one way ANOVA *F_2,13_* = 4.396, *p* = 0.0348, WS+VEH vs. WS+CNO p = 0.0414, WS+CNO vs. CON p = 0.3808). Representative Western Blot lanes depicting all treatment groups are presented in **Figure 3C** and full blots are presented in the Supplement (**Figure S3**). A separate group of rats were assessed for the behavioral effects of CNO in the absence of DREADD virus administration. These surgically naïve animals were assessed for burying behavior during acute WS with either VEH or CNO administered 1 hour prior, identical to Figure 3A except for the fact that there is no active DREADD virus in the LC. There was a significant effect of stress (**Figure 3D**, *F_1,23_* = 62.17, *p* < 0.0001) since WS exposed females treated with VEH displayed significantly higher levels of burying when compared to controls (p = 0.0002). Importantly, there was no effect of CNO on burying when the DREADD virus was absent, and WS+CNO-virus treated rats displayed significantly higher levels of burying when compared to controls (p = 0.0001), similar to the WS+VEH-virus group (p > 0.99). One animal from the WS+VEH-no virus group was eliminated from behavioral analysis due to a missing video file. Brains were collected from all animals in Experiment #2A, as well as all subsequent experiments, and virus localization to the LC was confirmed, with a representative image displaying the optimal atlas localization and LC virus spread in **Figure 3E**.

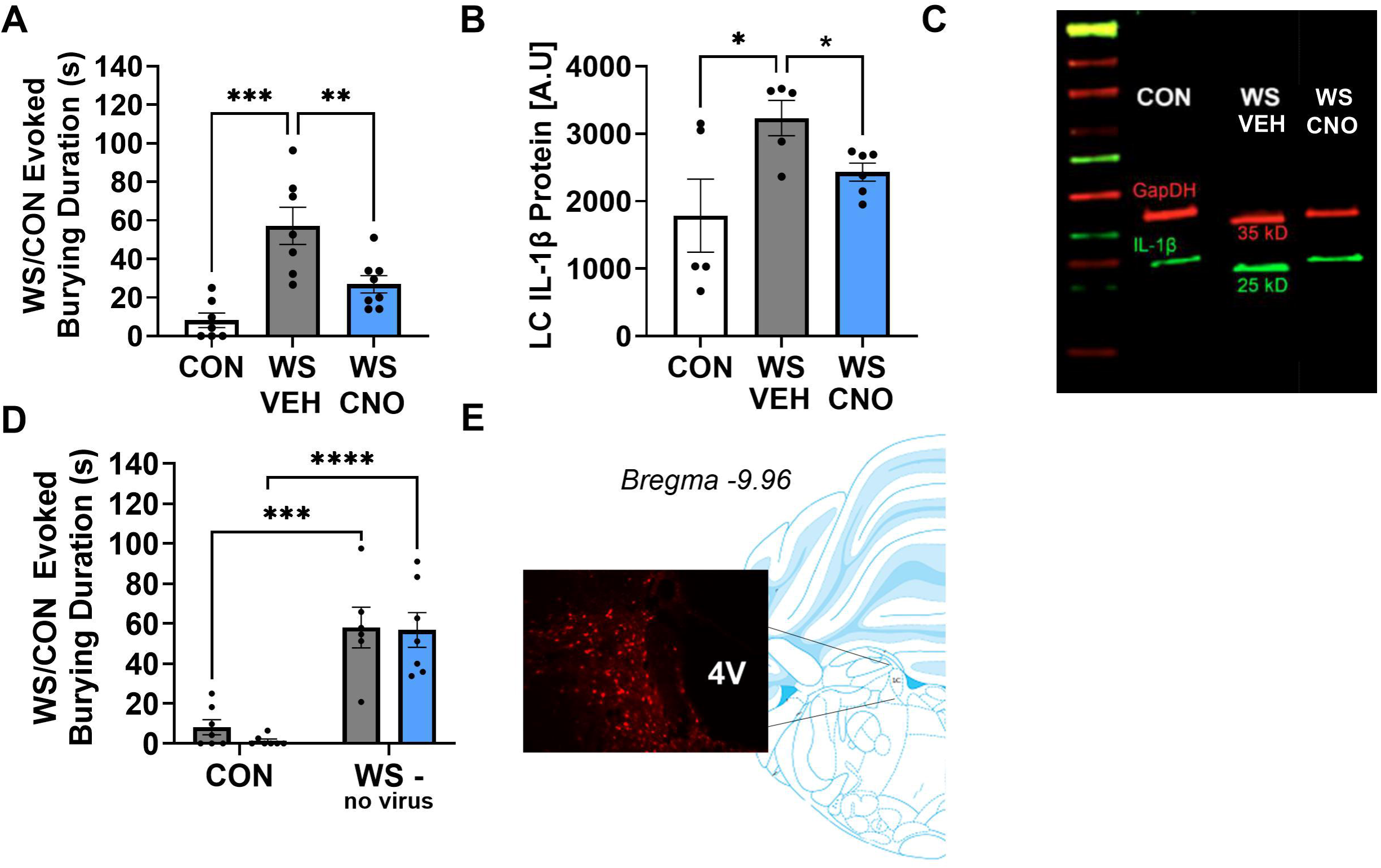

### Chemogenetic suppression of microglial reactivity prevents LPS-evoked microglial retraction

Experiment #3 was completed to determine the ability of microglial targeted DREADDs to affect microglia ramification characteristics as determined by computerized cell tracing and quantification. Sholl analysis of female rats treated with either VEH or LPS followed by VEH or CNO determined that systemic LPS administration resulted in a significantly lower number of intersections at all distances from the cell body when compared to VEH treated rats (**Figure 4C**, significant distance x treatment interaction, *F_24,108_* = 19.85, *p* < 0.0001) that was prevented with CNO administration (VEH+VEH vs. LPS+VEH p < 0.0001, LPS+VEH vs. LPS+CNO p < 0.0001, LPS+CNO). Additional measurements revealed an overall reactive cellular morphology in LPS treated rats that was not present with CNO treatment. LPS administration resulted in a significantly larger cell body area (**Figure 4D**, LPS effect *F_1,12_* = 37.70, *p* < 0.0001), lower number of projection endpoints (**Figure 4E**, LPS effect *F_1,12_* = 77.48, *p* < 0.0001), decreased total intersections (**Figure 4F**, LPS effect *F_1,12_* = 89.70, *p* < 0.0001), shorter projections (**Figure 4G**, LPS effect *F_1,12_* = 120.20, *p* < 0.0001), and a lower total projection length (**Figure 4H**, LPS effect *F_1,12_* = 79.57, *p* < 0.0001) when compared to controls, and all microglial ramification measures were all reversed with CNO treatment (CNO effect, cell body area *F_1,12_* = 40.92, *p* < 0.0001, number of endpoints *F_1,12_* = 52.46, *p* < 0.0001, total intersections *F_1,12_* = 45.53, *p* < 0.0001, projection length *F_1,12_* = 81.48, *p* < 0.0001, and total projection length *F_1,12_* = 39.92, *p* < 0.0001).

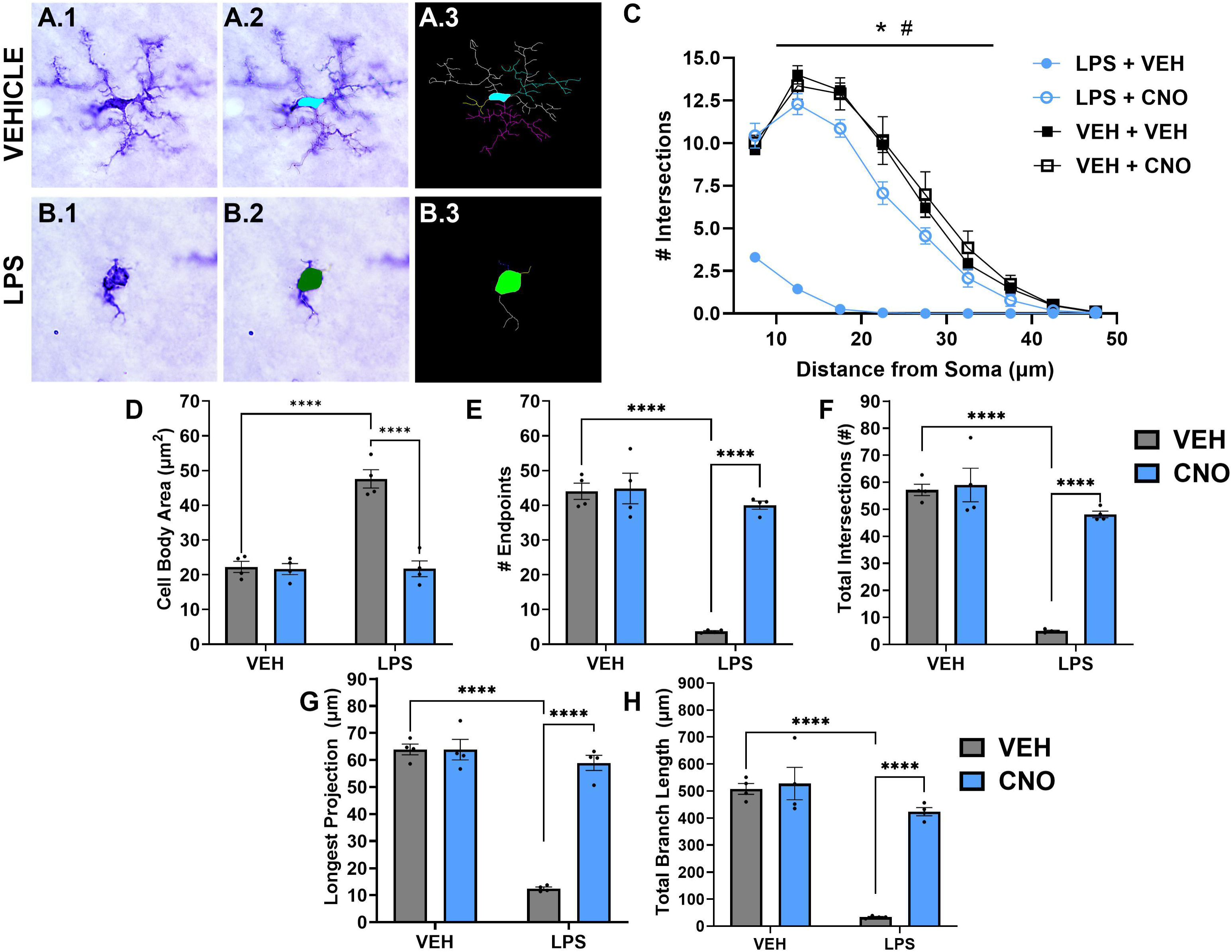

### DREADD activation prevents WS-induced changes in microglial ramification

To determine the impact of chemogenetic suppression of microglial reactivity on stress responsivity, rats were treated with VEH or CNO one hour prior to a single acute WS or control exposure followed by tissue collection for analysis of microglial ramification characteristics using Iba1 immunohistochemistry and computerized cell tracing and quantification. A subset of animals for this experiment did not receive surgery for infusion of the DREADDs virus to determine any off-target effects of CNO on microglial responses to stress. When assessing overall microglial ramification characteristics with a Sholl analysis, it was established that stressed females that received VEH as well as those that were treated with CNO in the absence of active virus displayed a fewer number of intersections across the distance from the cell body (**Figure 5C**, treatment group effect *F_5,162_* = 275.1, *p* < 0.0001). Further, the rats with active virus treated with CNO prior to stress exhibited an increase in cellular ramification, with a higher number of intersections across the distance from the cell body more similar to non-stressed controls (WS+VEH vs. WS+CNO p < 0.0001). When assessing additional microglial characteristics, WS resulted in a significant increase in cell body area (**Figure 5D**, *F_2,18_* = 27.22, *p* < 0.0001), decrease in the total number of branch endpoints (**Figure 5E**, *F_2,18_* = 161.50, *p* < 0.0001), decrease in the total number of intersections (**Figure 5F**, *F_2,18_* = 138.70, *p* < 0.0001), shorter projections (**Figure 5G**, *F_2,18_* = 122.20, *p* < 0.0001), and a lower total length of all projections (**Figure 5H**, *F_2,18_* = 126.90, *p* < 0.0001). Importantly, the increase in cell body size and decrease in projection complexity, number, and length were all prevented with CNO administration in rats with the active virus prior to stress exposure (CNO effect, cell body area *F_1,18_* = 14.50, *p* = 0.0013, number of endpoints *F_1,18_* = 11.53, *p* = 0.0032, total intersections *F_1,18_* = 14.55, *p* = 0.0013, longest projection *F_1,18_* = 10.54, *p* = 0.0045, and total projection length *F_1,18_* = 11.47, *p* = 0.0033) which did not occur in those without the active virus (WS+CNO vs. WS+CNO-no virus, cell body area *p* = 0.0007, number of endpoints *p* = 0.0002, total intersections *p* < 0.0001, longest projection *p* = 0.0012, and total projection length *p* = 0.0004).

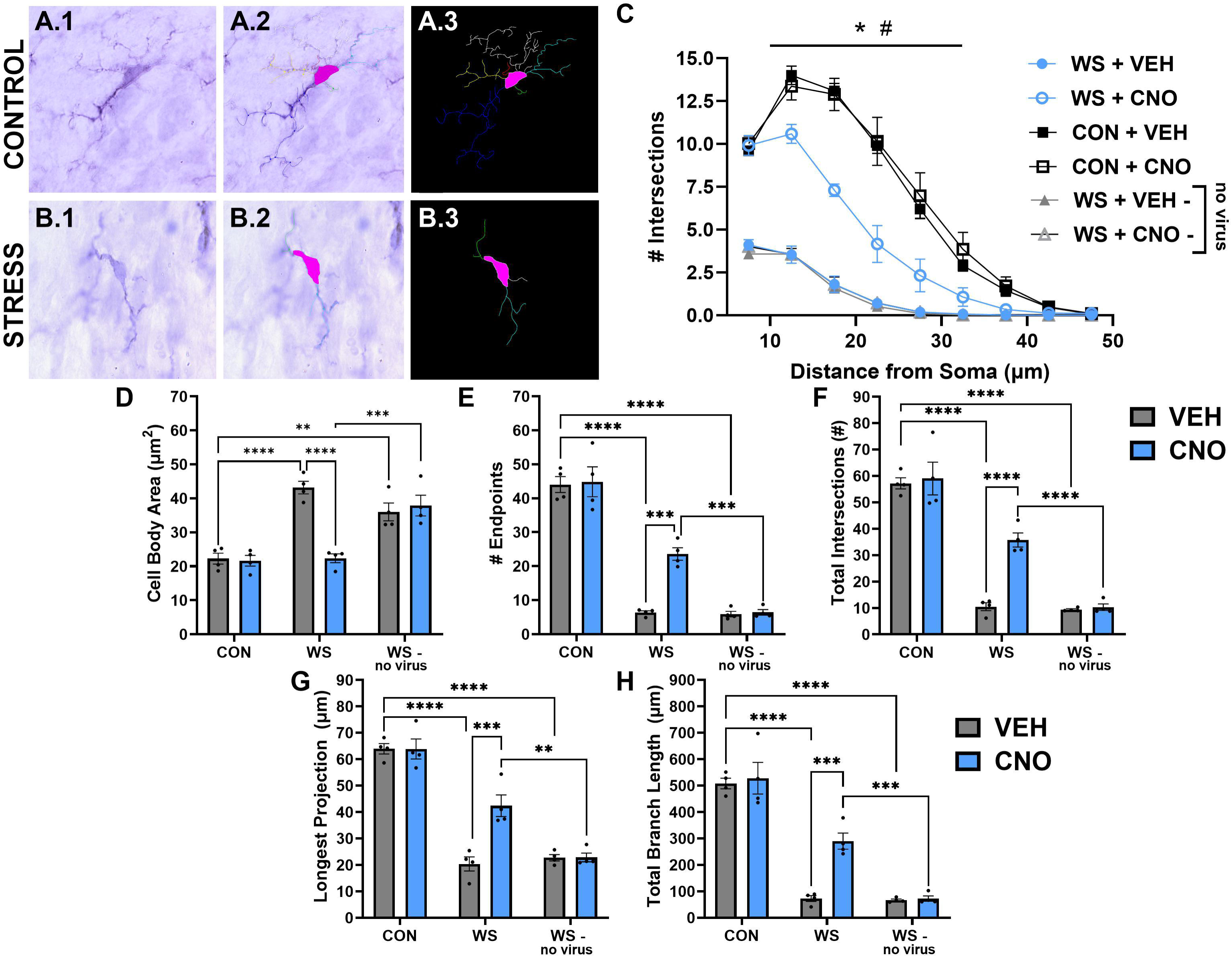

### Microglial inhibition prevents the behavioral effects of acute and repeated WS

A separate group of female rats for Experiment #4 were injected with the CD68-hM4Di-expressing virus selectively within the LC prior to assessment of behavioral tasks during acute and repeated stress and in response to the stress context. When first analyzing behaviors in response to acute WS, there was a significant effect of stress on burying behavior (**Figure 6A**, main effect of stress, *F_1,29_* = 18.30, *p* = 0.0002), replicating both our previous studies (Finnell et al., 2018; Pate et al., 2023; Smiley et al., 2023), additional studies using the WS model (Barbetti et al., 2023), and the results in **Figure 2A&B**. Additionally, CNO treatment one hour prior to the stressor diminished the burying response typically observed in females exposed to WS (**Figure 6A**, WS+VEH vs. WS+CNO *p* = 0.0048) which replicates the data presented in **Figure 2A**. Furthermore, when compared to non-stressed controls, WS exposed females exhibit a shorter latency to begin burying (**Figure 6B**, main effect of stress, *F_1,29_* = 41.43, *p* < 0.0001) and an increased duration of rearing (**Figure 6C**, *t_15_* = 1.548, *p* = 0.0712), both of which were attenuated by CNO treatment prior to stress (WS+VEH vs. WS+CNO, burying latency p = 0.0256, rearing duration p = 0.07). Freezing behavior was also assessed in response to stress, and no significant levels of freezing were observed in any treatment group (freezing duration, CON+VEH mean = 0 sec, CON+CNO mean = 0 sec, WS+VEH mean = 1.089 sec, WS+CNO mean = 0.038 sec). These acute effects were also observed in response to repeated stress. On WS Day #5, burying behavior was heightened in stressed females and significantly attenuated after CNO treatment prior to each stress exposure (**Figure 6D**, interaction, *F_1,28_* = 10.06, *p* = 0.0037; main effect of stress, *F_1,28_* = 19.74, *p* = 0.0001; main effect of drug, *F_1,28_* = 12.56, *p* = 0.0014; WS+VEH vs. WS+CNO *p* = 0.0002). While there was no longer a significant increase in the latency to begin burying in the CNO group due to an increased latency in the VEH treated animals (**Figure 6E**, main effect of stress *F_1,30_* = 74.46, *p* < 0.0001; WS+VEH vs. WS+CNO *p* = 0.3443), there was still a higher amount of rearing observed in stressed females that were treated with CNO (**Figure 6F**, main effect of stress *F_1,29_* = 51.94, *p* < 0.0001; WS+VEH vs. WS+CNO *p* = 0.0542). Additionally, there was no significant freezing observed across all groups in response to stress on Day 5 (freezing duration, CON+VEH mean = 0 sec, CON+CNO mean = 0 sec, WS+VEH mean = 0.863 sec, WS+CNO mean = 0.80 sec). Subsequently, when assessing stress cue-induced hypervigilance in the WS CXT, there was a significant interaction (*F_1,30_* = 19.69, *p* = 0.0001) between stress (main effect of stress, *F_1,30_* = 41.58, *p* < 0.0001) and drug (main effect of drug, *F_1,30_* = 20.05, *p* = 0.0001) in burying duration in response to the stress cues and environment (**Figure 6G**). Stressed females administered CNO displayed an increased latency to begin burying (**Figure 6H**, main effect of stress *F_1,30_* = 40.75, *p* < 0.0001; WS+VEH vs. WS+CNO *p* = 0.087) and no significant differences in the duration of rearing behavior (**Figure 6I**, main effect of stress *F_1,29_* = 22.37, *p* < 0.0001; WS+VEH vs. WS+CNO *p* = 0.337) when compared to vehicle treated rats in the WS CXT.

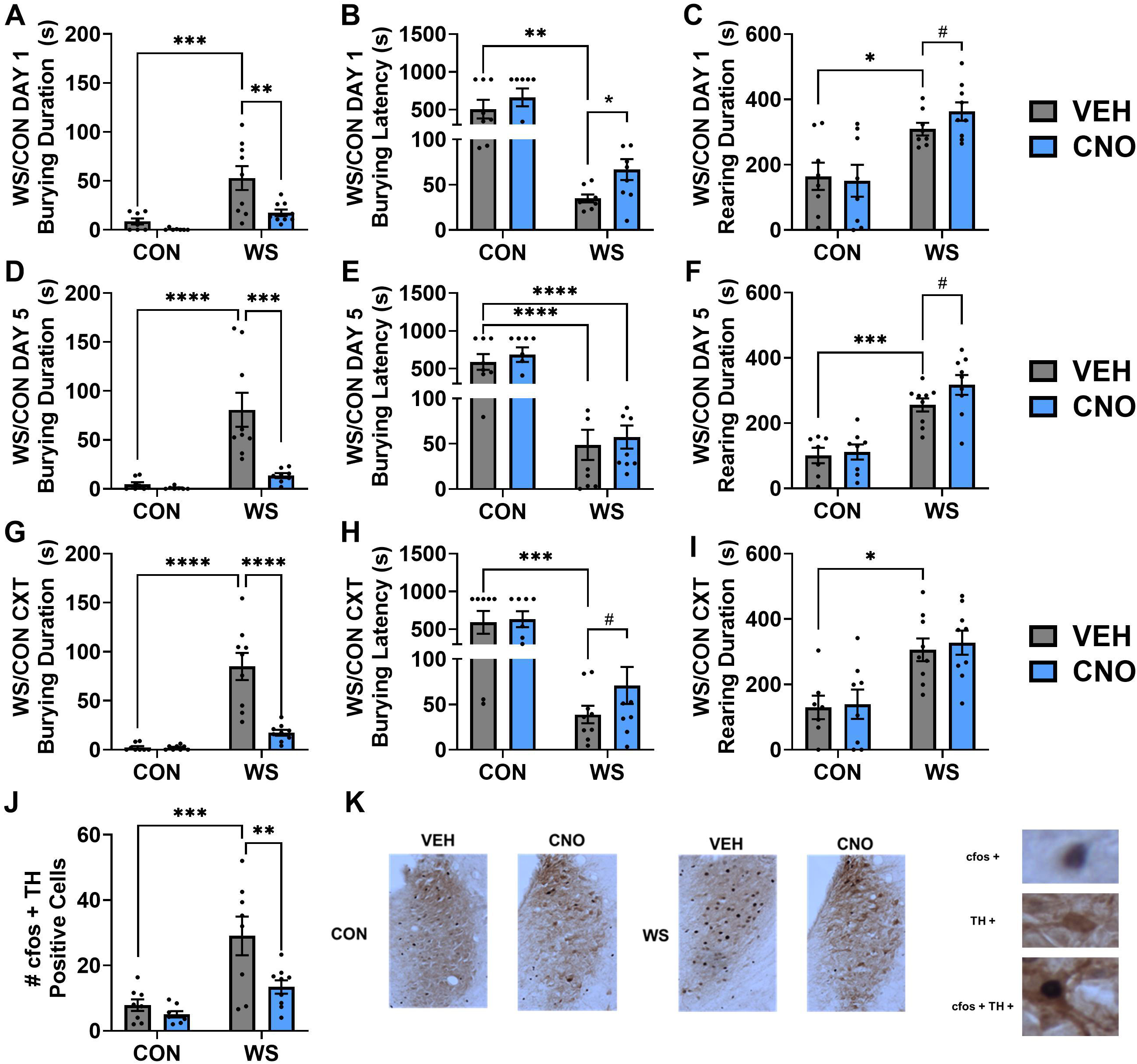

### WS CXT exposure increases LC activity which is prevented by a history of microglial suppression

Brain tissue was collected two hours following the start of WS/CON CXT for analysis of neuronal activity within the LC in rats with a history of CNO/VEH prior to repeated WS/CON. There was a significant effect of stress history (*F_1,29_* = 20.92, *p* < 0.0001) and drug treatment (*F_1,29_* = 8.126, *p* = 0.008) in the number of cfos positive TH labeled LC cells in response to the WS CXT (**Figure 6J**). Females exposed to WS+VEH exhibited a significantly higher level of neuronal activation in the LC in response to the stress cues when compared to controls (p = 0.0005) and to WS+CNO (p = 0.0084). Representative images of TH staining in the LC (brown) along with cfos positive cells (black) are shown for all treatment groups in **Figure 6K**.

## Discussion

These studies are among the first to utilize site specific microglial-targeted chemogenetics to manipulate neuroimmune activity within the brain and expand our recent evidence that microglia regulate LC-specific hypervigilant behavior resulting from witness stress in females (Pate et al., 2023). These experiments characterized the use of an inhibitory DREADD-expressing virus driven by the CD68 promoter within the LC and determined that this virus selectively colocalized with microglia, confirming published studies using these techniques in the spinal cord (Grace et al., 2016; Grace et al., 2018). Chemogenetic inhibition of microglia reduced WS-evoked inflammatory cytokine production, increased the complexity and ramification characteristics of these cells, and prevented WS-induced hypervigilant behaviors both in response to stress and stress cues. Notably, the DREADD-mediated reduction in stress-induced neuroimmune activity prevented the sensitization of hypervigilant behaviors in response to the WS CXT and was associated with a lower level of neuronal activity in the LC. Overall, these experiments have characterized the use of a CD68 promoter-driven hM4Di virus for targeting and manipulating microglial reactivity within the rat hindbrain and support the role of LC neuroimmune signaling in the acute and long-term impact of stress in females.

While chemogenetic techniques have been extensively utilized for their ability to impact neuronal signaling (Keifer et al., 2020; Urban & Roth, 2015), this method has only recently begun to be optimized for use in non-neuronal cell types (Bossuyt et al., 2023; Dheer et al., 2024; Parusel et al., 2023). AAVs, particularly AAV1, have been determined to effectively transduce microglial cells across the brain (Aschauer et al., 2013) and the reactivity of these cells, along with the resulting pro-inflammatory cytokine release, have also been shown to be regulated through GPCR-mediated mechanisms (Hayashi et al., 2011; Hoffmann et al., 2003). This virus has been developed to selectively colocalize with the Iba1 marker of microglia and not the astrocytic marker GFAP nor the neuronal NeuN and had a functional impact on the evoked expression of pro-inflammatory mediators IL-6, IL-1β, and TNFα (Grace et al., 2016; Grace et al., 2018). Importantly, viral transduction into microglia has been observed to have regional heterogeneity (Aschauer et al., 2013; Bossuyt et al., 2023), and these are the first experiments to observe a high level of microglial transduction within the LC. Further, species specific differences in microglial genetics, primarily the ∼6x higher levels of CD68 RNA observed in resting microglia in the rat versus the mouse (Lam et al., 2017), may influence differential rates of transfection between the species. Additional differences in the immune system have been observed between mice and rats, with species specific neuroimmune responses observed in response to multiple different types of inflammatory insults (Lively et al., 2018; Sroga et al., 2003). These studies identified a rat immune system as one that is more comparable to that of humans (Wildner, 2019) and, taken together, these data highlight the need to differentiate between the murine species when using novel tools to investigate the neuroimmune system. One further concern when using chemogenetic techniques is the risk for unintended effects of CNO outside of its function at the designed receptors. While higher doses of CNO, such as 10 mg/kg, produce off target effects and yield a high rate of conversion to clozapine, CNO at doses of 1 mg/kg, like those used in the present study, have been shown to have limited, if any, off target effects while plasma levels of clozapine are undetectable in rats (Manvich et al., 2018; Martinez et al., 2019). Importantly, these studies confirmed the lack of effect of CNO on behavior or microglial morphology shifts to alleviate these concerns.

Previous studies by our lab (Finnell et al., 2017; Pate et al., 2023; Wood et al., 2015) and others (Sanguino-Gómez et al., 2022; Schramm & Waisman, 2022; Sugama & Kakinuma, 2020) have highlighted the essential role of neuroimmune signaling in mediating CNS responses to stress (Biltz et al., 2022). While microglia are highly present throughout the brain, one region in which they are concentrated is the noradrenergic LC in which bidirectional regulation of norepinephrine and microglial cell reactivity plays a key role in the stress response (Iravani et al., 2014). Microglial responses can be regulated by adrenergic receptor signaling (Färber et al., 2005; Johnson et al., 2005) while, conversely, the release of proinflammatory cytokines leads to activation of the norepinephrine-producing LC cells (Borsody & Weiss, 2014). The concentration of cytokines such as IL-1β required to activate neuronal cells is ∼1,000 fold lower than that required by other cell types (Huang et al. 2011). Thus, even modest accumulations of IL-1β in the LC has the capacity to induce robust changes in neuronal activity as indicated by the current findings. Further, inflammatory mediated regulation of LC activity has been associated with hypervigilant and depressive-like behaviors (Kurosawa et al., 2016; Christoph Naegeli et al., 2018). Therefore, while these experiments determined that microglial reactivity plays a major role in the regulation of LC neuronal signaling and resulting hypervigilant burying behaviors, there is undoubtably further noradrenergic mechanisms involved in the regulation of this response. Additionally, while these studies focused on microglial signaling within the LC, peripheral immune signaling that occurs in response to stress exposure may also influence central immune activity. Recent studies have determined that there are time-, brain region-, and sex-specific effects on peripheral immune cell trafficking to the brain that may play a role in the stress response, with ∼10% of the immune cells of the hippocampus representing peripheral infiltration in response to acute learned helplessness paradigms (Medina-Rodriguez et al., 2023). Importantly, these results were not generalizable to females, chronic stress conditions, regions including the LC were not assessed, and this infiltration may result from local CNS immune signaling (Medina-Rodriguez et al., 2023). Therefore, while peripheral immune cell infiltration may play a role in microglial responses to stress, further experiments would be required to explore this mechanism in the female LC.

These experiments determined that microglial inhibition impacted behavioral responses to acute and repeated psychosocial stress, and this decrease in hypervigilant behavioral responding also occurred in response to subsequent exposure to the stress cues and environment. The inflammatory consequences of repeated stress have been shown to be sensitized with repeated stress exposure, a phenomenon often referred to as a “dual hit” hypothesis (Biltz et al., 2022; Smiley & Wood, 2022). The neuroimmune signaling and microglial reactivity that occurs in the brain as a result of social stress exposure leads to sensitization, whereby subthreshold stressor exposure in the future, i.e. exposure to the stress context, would therefore lead to an exaggerated inflammatory response (Biltz et al., 2022). Therefore, by preventing the inflammatory cascade that typically occurs in response to social stress (Biltz et al., 2022; Finnell et al., 2018; Liu et al., 2017; Pate et al., 2023; Wood et al., 2015) with chemogenetic inhibition of microglial activation, these results suggest that microglial priming and stress-induced sensitization of neuroimmune responding to subsequent stress and stress context exposures was blocked. Ultimately, we hypothesized that neuroimmune activity within the LC regulates hypervigilant behaviors through its effects on neuronal signaling, which was confirmed with cfos staining. Blocking microglial activation during stress was associated with lower levels of LC neuronal activation and reduced behavioral responding to stress cues. Future experiments utilizing *in vivo* electrophysiology to directly measure neuronal activity in response to microglial blockade will allow us to further probe the relationship between microglial and neuronal activity in the LC in the female stress response.

These studies identified a significant effect of LC manipulation on burying and rearing behavior in response to acute stress, repeated stress, and when presented with the stress cues and environment. Stressed females display a higher amount of burying along with less rearing behavior, which can be interpreted as an increase in compulsive anxiety-like behavior paralleled by a decrease in exploration. This data is supported by additional studies that have identified distinct LC projections to the central amygdala and medial prefrontal cortex that are responsible for increasing burying and decreasing exploration, respectively, in response to an external threat (Borodovitsyna et al., 2020). When assessing further behaviors that could be presented in response to stressful stimuli, no significant amounts of freezing were observed in any stressed females, emphasizing buying responses as the main anxiety-like behavior that is presented in response to a social stressor. While burying itself is not a translational measure, the ethological relevance of burying behavior in the rodent as a perseverative and repetitive behavior represents an important correlate to hypervigilant behaviors observed in humans with stress-related conditions such as PTSD (Mikics et al., 2008; Thomas et al., 2009). Further, burying behavior is associated with clinical signs of stress including increased stress hormone signaling and heart rate (Pate et al., 2023) and is highly dependent on the noradrenergic system (Liu et al., 2024), all which provide support for construct validity. Additionally, previous experiments from our lab have used translational measures of hypervigilance such as the acoustic startle task which can be completed similarly in rodents and humans. These studies have shown that our witness stress model evokes a higher startle response (Smiley et al., 2023) and further experiments would be required to determine the impact of LC immune reactivity on this translational measure of hypervigilance. While the LC is known to regulate hypervigilance and arousal in response to stressful experiences (Naegeli et al., 2018), this region also plays a role in conditional fear responding. Interestingly, LC projections are involved in hippocampal-dependent memory underlying the conditioned responses presented during associative fear learning (Wilmot et al., 2024). Furthermore, it is long known that norepinephrine signaling within the amygdala supports contextual fear learning (LaLumiere et al., 2003), a response that is escalated in females (Jasnow et al., 2006). Since these experiments involved multiple presentations of the stressor along with exposure to the stress cues, this model of psychosocial stress exposure may involve similar neural mechanisms to those required for contextual associative learning. Further, suppressing microglial reactivity in these studies may reduce LC activity and impair associative learning between stress exposure and stress cues. These questions serve as important future directions that would require further experimentation to establish the impact of direct microglial modulation in the LC on classical fear conditioning paradigms.

In conclusion, these experiments are among the first to site-specifically target microglia within the rat hindbrain for chemogenetic manipulation of neuroimmune signaling in response to stress. These studies confirmed our previous findings establishing LC microglia as a key player in the female stress response (Pate et al., 2023) and expanded the role of neuroimmune signaling to subsequent behavioral and neuronal responses to stress cues. Taken together, these experiments characterized the use of chemogenetics to alter microglial reactivity in the LC and support the role of LC neuroimmune signaling as a key mechanism in the female stress response. Subsequent experiments will further examine the functional relationship between immune activity, neuronal signaling and resulting NE output from the LC, as well as assessing sex differences, but the current results highlight the potential for targeting neuroimmune signaling in the treatment of stress-related comorbidities in females.

## Acknowledgements

Research reported in this publication was supported by the National Institute On Drug Abuse of the National Institutes of Health under Award Number F32-DA058380 (CES), National Institutes of Health (R01-MH113892, SKW), and Veterans Health Administration (I21 BX002664, SKW co-I and BX001374, SKW). Figures created with the help of BioRender.

## References

Aschauer, D. F., Kreuz, S., & Rumpel, S. (2013). Analysis of transduction efficiency, tropism and axonal transport of AAV serotypes 1, 2, 5, 6, 8 and 9 in the mouse brain. PLoS One, 8(9), e76310. 10.1371/journal.pone.0076310

Bangasser, D. A., & Valentino, R. J. (2014). Sex differences in stress-related psychiatric disorders: neurobiological perspectives. Front Neuroendocrinol, 35(3), 303–319. 10.1016/j.yfrne.2014.03.008

Barbetti, M., Vilella, R., Naponelli, V., Bilotti, I., Magistrati, M., Dallabona, C., Ielpo, D., Andolina, D., Sgoifo, A., Savi, M., & Carnevali, L. (2023). Repeated witness social stress causes cardiomyocyte contractile impairment and intracellular Ca2+ derangement in female rats. Physiol Behav, 271, 114339. 10.1016/j.physbeh.2023.114339

Bekhbat, M., & Neigh, G. N. (2018). Sex differences in the neuro-immune consequences of stress: Focus on depression and anxiety. Brain Behav Immun, 67, 1–12. 10.1016/j.bbi.2017.02.006

Benjet, C., Bromet, E., Karam, E. G., Kessler, R. C., McLaughlin, K. A., Ruscio, A. M., Shahly, V., Stein, D. J., Petukhova, M., Hill, E., Alonso, J., Atwoli, L., Bunting, B., Bruffaerts, R., Caldas-de-Almeida, J. M., de Girolamo, G., Florescu, S., Gureje, O., Huang, Y.,…Koenen, K. C. (2016). The epidemiology of traumatic event exposure worldwide: results from the World Mental Health Survey Consortium. Psychol Med, 46(2), 327–343. 10.1017/s0033291715001981

Biltz, R. G., Sawicki, C. M., Sheridan, J. F., & Godbout, J. P. (2022). The neuroimmunology of social-stress-induced sensitization. Nat Immunol, 23(11), 1527–1535. 10.1038/s41590-022-01321-z

Borodovitsyna, O., Duffy, B. C., Pickering, A. E., & Chandler, D. J. (2020). Anatomically and functionally distinct locus coeruleus efferents mediate opposing effects on anxiety-like behavior. Neurobiol Stress, 13, 100284. 10.1016/j.ynstr.2020.100284

Borsody, M. K., & Weiss, J. M. (2002). Alteration of locus coeruleus neuronal activity by interleukin-1 and the involvement of endogenous corticotropin-releasing hormone. Neuroimmunomodulation, 10(2), 101–121. 10.1159/000065186

Borsody, M. K., & Weiss, J. M. (2004). The effects of endogenous interleukin-1 bioactivity on locus coeruleus neurons in response to bacterial and viral substances. Brain Res, 1007(1), 39–56. 10.1016/j.brainres.2004.02.011

Borsody, M. K., & Weiss, J. M. (2014). Peripheral endotoxin causes long-lasting changes in locus coeruleus activity via IL-1 in the brain. Acta Neuropsychiatrica, 14(6), 303–321. 10.1034/j.1601-5215.2002.140605.x

Bossuyt, J., Van Den Herrewegen, Y., Nestor, L., Buckinx, A., De Bundel, D., & Smolders, I. (2023). Chemogenetic modulation of astrocytes and microglia: State-of-the-art and implications in neuroscience. Glia, 71(9), 2071–2095. 10.1002/glia.24390

Cohen, S., Janicki-Deverts, D., & Miller, G. E. (2007). Psychological Stress and Disease. JAMA, 298(14), 1685–1687. 10.1001/jama.298.14.1685

Davidson, J. R. (2000). Trauma: the impact of post-traumatic stress disorder. J Psychopharmacol, 14(2 Suppl 1), S5-12. 10.1177/02698811000142s102

Dheer, A., Bosco, D. B., Zheng, J., Wang, L., Zhao, S., Haruwaka, K., Yi, M. H., Barath, A., Tian, D. S., & Wu, L. J. (2024). Chemogenetic approaches reveal dual functions of microglia in seizures. Brain Behav Immun, 115, 406–418. 10.1016/j.bbi.2023.11.002

Engler, H., Benson, S., Wegner, A., Spreitzer, I., Schedlowski, M., & Elsenbruch, S. (2016). Men and women differ in inflammatory and neuroendocrine responses to endotoxin but not in the severity of sickness symptoms. Brain, Behavior, and Immunity, 52, 18–26. 10.1016/j.bbi.2015.08.013

Fan, Y., Chen, P., Li, Y., Cui, K., Noel, D. M., Cummins, E. D., Peterson, D. J., Brown, R. W., & Zhu, M. Y. (2014). Corticosterone administration up-regulated expression of norepinephrine transporter and dopamine β-hydroxylase in rat locus coeruleus and its terminal regions. J Neurochem, 128(3), 445–458. 10.1111/jnc.12459

Färber, K., Pannasch, U., & Kettenmann, H. (2005). Dopamine and noradrenaline control distinct functions in rodent microglial cells. Molecular and Cellular Neuroscience, 29(1), 128–138. 10.1016/j.mcn.2005.01.003

Finnell, Lombard, C. M., Melson, M. N., Singh, N. P., Nagarkatti, M., Nagarkatti, P., Fadel, J. R., Wood, C. S., & Wood, S. K. (2017). The protective effects of resveratrol on social stress-induced cytokine release and depressive-like behavior. Brain Behav Immun, 59, 147–157. 10.1016/j.bbi.2016.08.019

Finnell, Muniz, B. L., Padi, A. R., Lombard, C. M., Moffitt, C. M., Wood, C. S., Wilson, L. B., Reagan, L. P., Wilson, M. A., & Wood, S. K. (2018). Essential Role of Ovarian Hormones in Susceptibility to the Consequences of Witnessing Social Defeat in Female Rats. Biol Psychiatry, 84(5), 372–382. 10.1016/j.biopsych.2018.01.013

Frans, Ö., Rimmö, P.-A., Åberg, L., & Fredrikson, M. (2005). Trauma exposure and post-traumatic stress disorder in the general population. Acta Psychiatrica Scandinavica, 111(4), 291–290. 10.1111/j.1600-0447.2004.00463.x

Giustino, T. F., Ramanathan, K. R., Totty, M. S., Miles, O. W., & Maren, S. (2020). Locus Coeruleus Norepinephrine Drives Stress-Induced Increases in Basolateral Amygdala Firing and Impairs Extinction Learning. J Neurosci, 40(4), 907–916. 10.1523/jneurosci.1092-19.2019

Grace, P. M., Strand, K. A., Galer, E. L., Urban, D. J., Wang, X., Baratta, M. V., Fabisiak, T. J., Anderson, N. D., Cheng, K., Greene, L. I., Berkelhammer, D., Zhang, Y., Ellis, A. L., Yin, H. H., Campeau, S., Rice, K. C., Roth, B. L., Maier, S. F., & Watkins, L. R. (2016). Morphine paradoxically prolongs neuropathic pain in rats by amplifying spinal NLRP3 inflammasome activation. Proc Natl Acad Sci U S A, 113(24), E3441–3450. 10.1073/pnas.1602070113

Grace, P. M., Wang, X., Strand, K. A., Baratta, M. V., Zhang, Y., Galer, E. L., Yin, H., Maier, S. F., & Watkins, L. R. (2018). DREADDed microglia in pain: Implications for spinal inflammatory signaling in male rats. Exp Neurol, 304, 125–131. 10.1016/j.expneurol.2018.03.005

Gu, C., Chen, Y., Chen, Y., Liu, C. F., Zhu, Z., & Wang, M. (2021). Role of G Protein-Coupled Receptors in Microglial Activation: Implication in Parkinson’s Disease. Front Aging Neurosci, 13, 768156. 10.3389/fnagi.2021.768156

Hänsel, A., Hong, S., Cámara, R. J. A., & von Känel, R. (2010). Inflammation as a psychophysiological biomarker in chronic psychosocial stress. Neuroscience & Biobehavioral Reviews, 35(1), 115–121. 10.1016/j.neubiorev.2009.12.012

Haque, M. E., Kim, I.-S., Jakaria, M., Akther, M., & Choi, D.-K. (2018). Importance of GPCR-Mediated Microglial Activation in Alzheimer’s Disease [Review]. Front Cell Neurosci, 12. 10.3389/fncel.2018.00258

Hayashi, Y., Kawaji, K., Sun, L., Zhang, X., Koyano, K., Yokoyama, T., Kohsaka, S., Inoue, K., & Nakanishi, H. (2011). Microglial Ca(2+)-activated K(+) channels are possible molecular targets for the analgesic effects of S-ketamine on neuropathic pain. J Neurosci, 31(48), 17370–17382. 10.1523/jneurosci.4152-11.2011

Hoffmann, A., Kann, O., Ohlemeyer, C., Hanisch, U. K., & Kettenmann, H. (2003). Elevation of basal intracellular calcium as a central element in the activation of brain macrophages (microglia): suppression of receptor-evoked calcium signaling and control of release function. J Neurosci, 23(11), 4410–4419. 10.1523/jneurosci.23-11-04410.2003

Iravani, M. M., Sadeghian, M., Rose, S., & Jenner, P. (2014). Loss of locus coeruleus noradrenergic neurons alters the inflammatory response to LPS in substantia nigra but does not affect nigral cell loss. Journal of Neural Transmission, 121(12), 1493–1505. 10.1007/s00702-014-1223-1

Irish, L. A., Fischer, B., Fallon, W., Spoonster, E., Sledjeski, E. M., & Delahanty, D. L. (2011). Gender differences in PTSD symptoms: an exploration of peritraumatic mechanisms. J Anxiety Disord, 25(2), 209–216. 10.1016/j.janxdis.2010.09.004

Jasnow, A. M., Schulkin, J., & Pfaff, D. W. (2006). Estrogen facilitates fear conditioning and increases corticotropin-releasing hormone mRNA expression in the central amygdala in female mice. Horm Behav, 49(2), 197–205. 10.1016/j.yhbeh.2005.06.005

Johnson, J. D., Campisi, J., Sharkey, C. M., Kennedy, S. L., Nickerson, M., Greenwood, B. N., & Fleshner, M. (2005). Catecholamines mediate stress-induced increases in peripheral and central inflammatory cytokines. Neuroscience, 135(4), 1295–1307. 10.1016/j.neuroscience.2005.06.090

Kedia, S., & Chattarji, S. (2014). Marble burying as a test of the delayed anxiogenic effects of acute immobilisation stress in mice. Journal of Neuroscience Methods, 233, 150–154. 10.1016/j.jneumeth.2014.06.012

Keifer, O., Kambara, K., Lau, A., Makinson, S., & Bertrand, D. (2020). Chemogenetics a robust approach to pharmacology and gene therapy. Biochemical Pharmacology, 175, 113889. 10.1016/j.bcp.2020.113889

Kettenmann, H., Hanisch, U. K., Noda, M., & Verkhratsky, A. (2011). Physiology of microglia. Physiol Rev, 91(2), 461–553. 10.1152/physrev.00011.2010

Kimble, M., Boxwala, M., Bean, W., Maletsky, K., Halper, J., Spollen, K., & Fleming, K. (2014). The impact of hypervigilance: evidence for a forward feedback loop. J Anxiety Disord, 28(2), 241–245. 10.1016/j.janxdis.2013.12.006

Krystal, J. H., Abdallah, C. G., Pietrzak, R. H., Averill, L. A., Harpaz-Rotem, I., Levy, I., Kelmendi, B., & Southwick, S. M. (2018). Locus Coeruleus Hyperactivity in Posttraumatic Stress Disorder: Answers and Questions. Biological Psychiatry, 83(3), 197–199. 10.1016/j.biopsych.2017.09.027

Kurosawa, N., Shimizu, K., & Seki, K. (2016). The development of depression-like behavior is consolidated by IL-6-induced activation of locus coeruleus neurons and IL-1β-induced elevated leptin levels in mice. Psychopharmacology, 233(9), 1725–1737. 10.1007/s00213-015-4084-x

LaLumiere, R. T., Buen, T. V., & McGaugh, J. L. (2003). Post-training intra-basolateral amygdala infusions of norepinephrine enhance consolidation of memory for contextual fear conditioning. J Neurosci, 23(17), 6754–6758. 10.1523/jneurosci.23-17-06754.2003

Lam, D., Lively, S., & Schlichter, L. C. (2017). Responses of rat and mouse primary microglia to pro- and anti-inflammatory stimuli: molecular profiles, K(+) channels and migration. J Neuroinflammation, 14(1), 166. 10.1186/s12974-017-0941-3

Levesque, P., Desmeules, C., Béchard, L., Huot-Lavoie, M., Demers, M.-F., Roy, M.-A., & Deslauriers, J. (2023). Sex-specific immune mechanisms in PTSD symptomatology and risk: A translational overview and perspectives. Brain Research Bulletin, 195, 120–129. 10.1016/j.brainresbull.2023.02.013

Liberzon, I., & Abelson, J. L. (2016). Context Processing and the Neurobiology of Post-Traumatic Stress Disorder. Neuron, 92(1), 14–30. 10.1016/j.neuron.2016.09.039

Liu, J., Lustberg, D. J., Galvez, A., Liles, L. C., McCann, K. E., & Weinshenker, D. (2024). Genetic disruption of dopamine β-hydroxylase dysregulates innate responses to predator odor in mice. Neurobiol Stress, 29, 100612. 10.1016/j.ynstr.2024.100612

Liu, Y. Z., Wang, Y. X., & Jiang, C. L. (2017). Inflammation: The Common Pathway of Stress-Related Diseases. Front Hum Neurosci, 11, 316. 10.3389/fnhum.2017.00316

Lively, S., Lam, D., Wong, R., & Schlichter, L. C. (2018). Comparing Effects of Transforming Growth Factor β1 on Microglia From Rat and Mouse: Transcriptional Profiles and Potassium Channels. Front Cell Neurosci, 12, 115. 10.3389/fncel.2018.00115

Manvich, D. F., Webster, K. A., Foster, S. L., Farrell, M. S., Ritchie, J. C., Porter, J. H., & Weinshenker, D. (2018). The DREADD agonist clozapine N-oxide (CNO) is reverse-metabolized to clozapine and produces clozapine-like interoceptive stimulus effects in rats and mice. Scientific Reports, 8(1), 3840. 10.1038/s41598-018-22116-z

Marshall, G. N., Schell, T. L., Glynn, S. M., & Shetty, V. (2006). The role of hyperarousal in the manifestation of posttraumatic psychological distress following injury. J Abnorm Psychol, 115(3), 624–628. 10.1037/0021-843x.115.3.624

Martinez, V. K., Saldana-Morales, F., Sun, J. J., Zhu, P. J., Costa-Mattioli, M., & Ray, R. S. (2019). Off-Target Effects of Clozapine-N-Oxide on the Chemosensory Reflex Are Masked by High Stress Levels. Front Physiol, 10, 521. 10.3389/fphys.2019.00521

McCall, J. G., Al-Hasani, R., Siuda, E. R., Hong, D. Y., Norris, A. J., Ford, C. P., & Bruchas, M. R. (2015). CRH Engagement of the Locus Coeruleus Noradrenergic System Mediates Stress-Induced Anxiety. Neuron, 87(3), 605–620. 10.1016/j.neuron.2015.07.002

McCall, J. G., Siuda, E. R., Bhatti, D. L., Lawson, L. A., McElligott, Z. A., Stuber, G. D., & Bruchas, M. R. (2017). Locus coeruleus to basolateral amygdala noradrenergic projections promote anxiety-like behavior. Elife, 6. 10.7554/eLife.18247

Medina-Rodriguez, E. M., Han, D., Lowell, J., & Beurel, E. (2023). Stress promotes the infiltration of peripheral immune cells to the brain. *Brain*, Behavior, and Immunity, 111, 412–423. 10.1016/j.bbi.2023.05.003

Miao, X. R., Chen, Q. B., Wei, K., Tao, K. M., & Lu, Z. J. (2018). Posttraumatic stress disorder: from diagnosis to prevention. Mil Med Res, 5(1), 32. 10.1186/s40779-018-0179-0

Mikics, E., Baranyi, J., & Haller, J. (2008). Rats exposed to traumatic stress bury unfamiliar objects--a novel measure of hyper-vigilance in PTSD models? Physiol Behav, 94(3), 341–348. 10.1016/j.physbeh.2008.01.023

Mizoguchi, Y., & Monji, A. (2017). Microglial Intracellular Ca(2+) Signaling in Synaptic Development and its Alterations in Neurodevelopmental Disorders. Front Cell Neurosci, 11, 69. 10.3389/fncel.2017.00069

Moieni, M., Irwin, M. R., Jevtic, I., Olmstead, R., Breen, E. C., & Eisenberger, N. I. (2015). Sex differences in depressive and socioemotional responses to an inflammatory challenge: implications for sex differences in depression. Neuropsychopharmacology, 40(7), 1709–1716. 10.1038/npp.2015.17

Naegeli, Zeffiro, T., Piccirelli, M., Jaillard, A., Weilenmann, A., Hassanpour, K., Schick, M., Rufer, M., Orr, S. P., & Mueller-Pfeiffer, C. (2018). Locus Coeruleus Activity Mediates Hyperresponsiveness in Posttraumatic Stress Disorder. Biol Psychiatry, 83(3), 254–262. 10.1016/j.biopsych.2017.08.021

Naegeli, C., Zeffiro, T., Piccirelli, M., Jaillard, A., Weilenmann, A., Hassanpour, K., Schick, M., Rufer, M., Orr, S. P., & Mueller-Pfeiffer, C. (2018). Locus Coeruleus Activity Mediates Hyperresponsiveness in Posttraumatic Stress Disorder. Biol Psychiatry, 83(3), 254–262. 10.1016/j.biopsych.2017.08.021

Naegeli, C., Zeffiro, T., Piccirelli, M., Jaillard, A., Weilenmann, A., Hassanpour, K., Schick, M., Rufer, M., Orr, S. P., & Mueller-Pfeiffer, C. (2018). Locus Coeruleus Activity Mediates Hyperresponsiveness in Posttraumatic Stress Disorder. Biological Psychiatry, 83(3), 254–262. 10.1016/j.biopsych.2017.08.021

O’Donnell, T., Hegadoren, K. M., & Coupland, N. C. (2004). Noradrenergic mechanisms in the pathophysiology of post-traumatic stress disorder. Neuropsychobiology, 50(4), 273–283. 10.1159/000080952

Parusel, S., Yi, M. H., Hunt, C. L., & Wu, L. J. (2023). Chemogenetic and Optogenetic Manipulations of Microglia in Chronic Pain. Neurosci Bull, 39(3), 368–378. 10.1007/s12264-022-00937-3

Pate, B. S., Bouknight, S. J., Harrington, E. N., Mott, S. E., Augenblick, L. M., Smiley, C. E., Morgan, C. G., Calatayud, B. M., Martínez-Muñiz, G. A., Thayer, J. F., & Wood, S. K. (2023). Site-Specific knockdown of microglia in the locus coeruleus regulates hypervigilant responses to social stress in female rats. *Brain*, Behavior, and Immunity, 109, 190–203. 10.1016/j.bbi.2023.01.011

Piña, R., Santos-Díaz, A. I., Orta-Salazar, E., Aguilar-Vazquez, A. R., Mantellero, C. A., Acosta-Galeana, I., Estrada-Mondragon, A., Prior-Gonzalez, M., Martinez-Cruz, J. I., & Rosas-Arellano, A. (2022). Ten Approaches That Improve Immunostaining: A Review of the Latest Advances for the Optimization of Immunofluorescence. Int J Mol Sci, 23(3). 10.3390/ijms23031426

Ravi, M., Miller, A. H., & Michopoulos, V. (2021). The immunology of stress and the impact of inflammation on the brain and behaviour. BJPsych Advances, 27(3), 158–165. 10.1192/bja.2020.82

Sadeghi, M., McDonald, A. D., & Sasangohar, F. (2022). Posttraumatic stress disorder hyperarousal event detection using smartwatch physiological and activity data. PLoS One, 17(5), e0267749. 10.1371/journal.pone.0267749

Sanguino-Gómez, J., Buurstede, J. C., Abiega, O., Fitzsimons, C. P., Lucassen, P. J., Eggen, B. J. L., Lesuis, S. L., Meijer, O. C., & Krugers, H. J. (2022). An emerging role for microglia in stress-effects on memory. Eur J Neurosci, 55(9-10), 2491–2518. 10.1111/ejn.15188

Schell, T. L., Marshall, G. N., & Jaycox, L. H. (2004). All Symptoms Are Not Created Equal: The Prominent Role of Hyperarousal in the Natural Course of Posttraumatic Psychological Distress. Journal of Abnormal Psychology, 113, 189–197. 10.1037/0021-843X.113.2.189

Schramm, E., & Waisman, A. (2022). Microglia as Central Protagonists in the Chronic Stress Response. Neurol Neuroimmunol Neuroinflamm, 9(6). 10.1212/nxi.0000000000200023

Smiley, C. E., Pate, B. S., Bouknight, S. J., Francis, M. J., Nowicki, A. V., Harrington, E. N., & Wood, S. K. (2023). Estrogen receptor beta in the central amygdala regulates the deleterious behavioral and neuronal consequences of repeated social stress in female rats. Neurobiol Stress, 23, 100531. 10.1016/j.ynstr.2023.100531

Smiley, C. E., & Wood, S. K. (2022). Stress- and drug-induced neuroimmune signaling as a therapeutic target for comorbid anxiety and substance use disorders. Pharmacology & Therapeutics, 239, 108212. 10.1016/j.pharmthera.2022.108212

Sroga, J. M., Jones, T. B., Kigerl, K. A., McGaughy, V. M., & Popovich, P. G. (2003). Rats and mice exhibit distinct inflammatory reactions after spinal cord injury. J Comp Neurol, 462(2), 223–240. 10.1002/cne.10736

Sugama, S., & Kakinuma, Y. (2020). Stress and brain immunity: Microglial homeostasis through hypothalamus-pituitary-adrenal gland axis and sympathetic nervous system. Brain Behav Immun Health, 7, 100111. 10.1016/j.bbih.2020.100111

Thomas, A., Burant, A., Bui, N., Graham, D., Yuva-Paylor, L. A., & Paylor, R. (2009). Marble burying reflects a repetitive and perseverative behavior more than novelty-induced anxiety. Psychopharmacology (Berl), 204(2), 361–373. 10.1007/s00213-009-1466-y

Urban, D. J., & Roth, B. L. (2015). DREADDs (Designer Receptors Exclusively Activated by Designer Drugs): Chemogenetic Tools with Therapeutic Utility. Annual Review of Pharmacology and Toxicology, 55(1), 399–417. 10.1146/annurev-pharmtox-010814-124803

Wildner, G. (2019). Are rats more human than mice? Immunobiology, 224(1), 172–176. 10.1016/j.imbio.2018.09.002

Wilmot, J. H., Diniz, C., Crestani, A. P., Puhger, K. R., Roshgadol, J., Tian, L., & Wiltgen, B. J. (2024). Phasic locus coeruleus activity enhances trace fear conditioning by increasing dopamine release in the hippocampus. Elife, 12. 10.7554/eLife.91465

Wood, S. K., Wood, C. S., Lombard, C. M., Lee, C. S., Zhang, X. Y., Finnell, J. E., & Valentino, R. J. (2015). Inflammatory Factors Mediate Vulnerability to a Social Stress-Induced Depressive-like Phenotype in Passive Coping Rats. Biol Psychiatry, 78(1), 38–48. 10.1016/j.biopsych.2014.10.026

Yi, M. H., Liu, Y. U., Liu, K., Chen, T., Bosco, D. B., Zheng, J., Xie, M., Zhou, L., Qu, W., & Wu, L. J. (2021). Chemogenetic manipulation of microglia inhibits neuroinflammation and neuropathic pain in mice. Brain Behav Immun, 92, 78–89. 10.1016/j.bbi.2020.11.030

