## Supplemental Figures for "The functional role of locus coeruleus microglia in the female stress response"

**Supplementary Materials for: The role of locus coeruleus neuroimmune signaling in the response to social stress in female rats**

**Supplementary Figure 1.** Virus placement localization within the LC for all treatment groups.

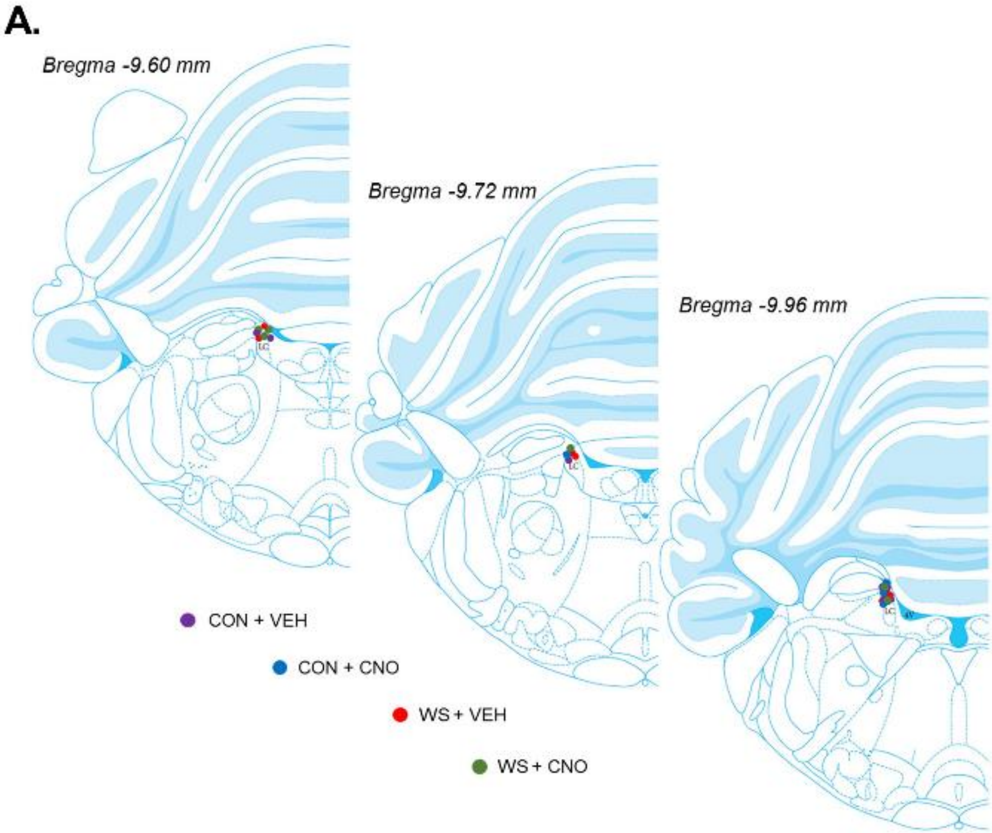


**Supplementary Figure 2.** The location of micropunches taken to include the locus coeruleus.

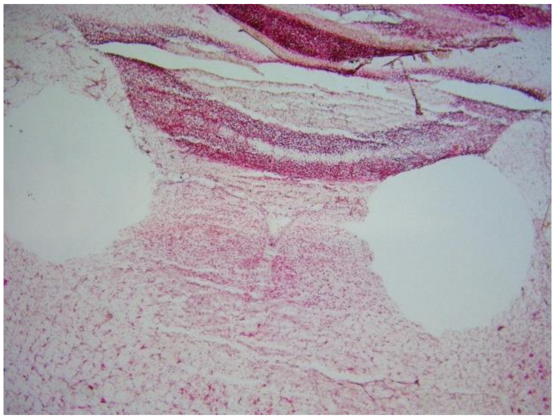

**Supplementary Figure 3.** Full Western Blot images for Figure 3B with GapDH (red) and IL-1β (green).

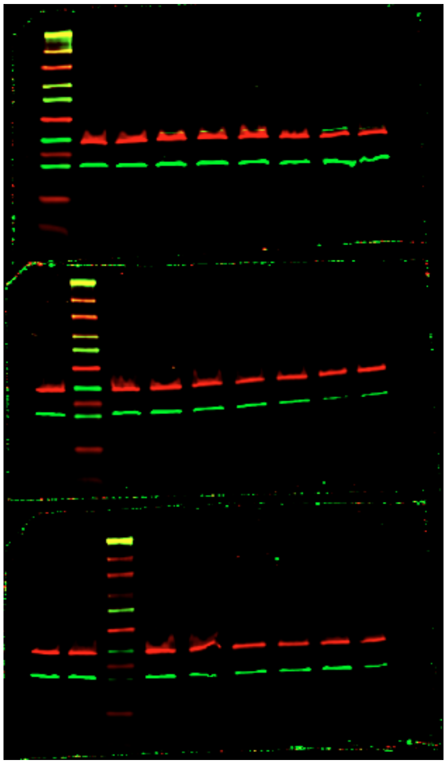
